# Human mediated dispersal of cats in the Neolithic Central Europe

**DOI:** 10.1101/259143

**Authors:** Mateusz Baca, Danijela Popović, Hanna Panagiotopoulou, Adrian Marciszak, Magdalena Krajcarz, Maciej T. Krajcarz, Daniel Makowiecki, Piotr Węgleński, Adam Nadachowski

## Abstract

Archaeological and genetic evidence suggest that all domestic cats derive from the Near Eastern wildcat (*Felis silvestris lybica*) and were domesticated twice, first in the Near East around 10 000 years ago and for the second time in Ancient Egypt ca. 3 500 years ago. The spread of the domesticated form in Europe occurred much later, primarily mediated by Greek and Phoenician traders and afterwards by Romans who introduced cats to Western and Central Europe around 2 000 years ago. We investigated mtDNA of Holocene *Felis* remains and provide evidence of an unexpectedly early presence of cats bearing the Near Eastern wildcat mtDNA haplotypes in Central Europe, being ahead of Roman Period by over 2 000 years. The appearance of the Near Eastern wildcats in Central Europe coincide with the peak of Neolithic settlement density, moreover most of those cats belonged to the same mtDNA lineages as those domesticated in the Near East. Thus, although we cannot fully exclude that the Near Eastern wildcats appeared in Central Europe as a result of introgression with European wildcat, our findings strongly support the hypothesis that the Near Eastern wildcats spread across Europe together with the first farmers, perhaps as commensal animals. We also found that cats dated to the Neolithic period belonged to different mtDNA lineages than those brought to Central Europe in Roman times, this supports the hypothesis that the gene pool of contemporary European domestic cats might have been established from two different source populations that contributed in different periods.

## Introduction

Latest research on the wildcat *Felis silvestris* phylogeny resulted in distinction of five subspecies rank groups, corresponding to their geographical distribution (Driscoll *et al.*, 2007, 2009): European wildcat, Southern African wildcat, Central Asian wildcat, Near Eastern wildcat and Chinese desert cat. Today, the European range of *F. silvestris* includes two subspecies. The European wildcat *(F. s. silvestris)* represents the only native form in most of the region. This animal was formerly widely distributed in Europe, except Fennoscandia (Yamaguchi *et al.*, 2015), however has become extinct in many areas mainly due to hunting and habitat loses. The second subspecies, domestic cat (*F. s.catus*), is of anthropogenic origin. The history of cat domestication was reconstructed with use of preserved written sources, art objects and archaeozoological material and significantly supported during last years with genetic studies. According to genetic data, domestic cats do not descend from European wildcat, although European wildcat and domestic cat may share territory and crossbreed as interfertile taxa. The common ancestor to all modern domestic cats was the Near Eastern wildcat, *F. s. lybica* (Driscoll *et al.*, 2007), domesticated in the Near East during Neolithic (Vigne *et al.*, 2004; Driscoll *et al.*, 2009; Faure and Kitchener, 2009; Ottoni *et al.*, 2017). The descendants of the domesticated Near Eastern wildcats were later spread across the world along with civilization expansion. Recently, analysis of mtDNA from more than 200 *Felis* remains revealed that cats were domesticated from at least two different local populations of the Near Eastern wildcats, for the first time in the Near East and for the second in Ancient Egypt (Ottoni *et al.*, 2017).

According to the current knowledge, the domestic cat did not occur in Central Europe prior to Roman Period (Benecke, 1994; Clutton-Brock, 1999; Driscoll *et al.*, 2009; Faure and Kitchener, 2009; Krajcarz *et al.*, 2016), however the chronology of the Near Eastern wildcat introduction to different regions of Europe is still weakly understood. The archaeozoological and paleontological records are poor and direct chronometric data and ancient DNA analyses of fossil cats are still rare.

The preliminary study about the history of domestic cats in Poland (Krajcarz *et al.*, 2016) revealed no presence of domesticated forms in archaeological contexts before 1^st^ century AD. Since that study was focused on archaeozoological material and did not include cat remains from non-human related sites, there was a risk of overlooking the natural or civilization related expansions of cats from the Near East. Here, we extended the prior survey to fossil Holocene cat’s remains recovered from outside the archaeological contexts.

## Materials and Methods

We analysed bone fragments of 36 individuals from 19 sites in Poland (Supplementary Table S1) that were provisionally classified as *Felis* sp. or *Felis silvestris*. This include six specimens excavated from archaeological contexts for which partial mtDNA ND5 sequence was already published (Krajcarz *et al.*, 2016). Sample handling and DNA extraction were performed in a laboratory dedicated to ancient DNA analyses in the Laboratory of Paleogenetics and Conservation Genetics, Centre of New Technologies at the University of Warsaw. Strict contamination precautions were undertaken during all steps of the experimental procedure. Prior to DNA extraction, each sample was washed with bleach solution (6%w/v sodium hypochlorite), rinsed with double distilled water, UV-irradiated (245 nm) for 20 minutes on each side and pulverized in cryogenic mill (SPEX CentriPrep, Stanmore, UK). DNA extraction was performed using modified silica column based method optimized to retrieve short DNA fragments (Dabney *et al.*, 2013). Samples were processed in batches of 16 with a negative control included in each batch. First we screened all samples for DNA preservation by amplification of a short fragment of mitochondrial ND5 gene. Thirty-three samples yielded DNA sequence that allow initial species assignation (Supplementary Table 1). To obtain longer fragment of the mtDNA sequences we used a targeted enrichment approach. For the hybridization experiment we choose 20 samples, 12 that yielded *F. s. lybica/catus*, and eight that yielded *F.s. silvestris* haplotypes during initial screening. Those samples were either already radiocarbon dated or there was enough bone left to perform dating. DNA extracts were converted into double-indexed sequencing libraries following modified protocol of Kircher and Meyer (2010). To minimize sample cross-talk during sequencing, beside double-indexing, we used adapters containing 7 bp long barcodes (Rohland *et al.*, 2014). We targeted a 6 kb fragment of mtDNA genome spanning position from 11 487 to 925. Hybridization bait was produced from the DNA of a contemporary domestic cat. DNA from swab was extracted with DNeasy Blood & Tissue Kit (Qiagen), and then the desired mtDNA fragment was amplified with three primer pairs. PCR products were sonicated to the length of around 200 bp with Covaris S220 and converted into bait following the protocol of Maricic *et al.* (2010). Hybridization was carried on pools of up to five libraries. We performed two rounds of hybridisation for 21h each following the protocol proposed by Horn (2012). Libraries were amplified for 19 cycles after the first and for 17 cycles after the second round. Enriched libraries were quantified with qPCR (Illumina Library Quantification kit, KAPA), pooled in equimolar ratios and sequenced with other libraries on NextSeq or on MiSeq platform (Illumina) in the 2 x 75 bp or 2 x 150 modes, respectively. Libraries produced from extraction negative controls were pooled, hybridized and sequenced as other libraries.

Sequencing reads were demultiplexed using Bcl2fastq, reads containing appropriate barcode were filtered with Sabre script, and then AdapterRemoval v. 2 (Lindgreen, 2012) was used to collapse overlapping reads. Reads were mapped to cat reference mtDNA sequence using Bwa (Li and Durbin, 2010), only reads with mapping quality over 30 and longer than 30 bp were retained. Duplicates were removed; variants and consensus sequences were called using Samtools and Bcftools (Li *et al.*, 2009). We called only positions with minimum 2 x coverage. Each bam alignment was inspected manually in Tablet (Milne *et al.*, 2013). Endogenous ancient DNA molecules typically exhibit excess of deaminated cytosine towards the ends of molecules; we used MapDamage v.2 (Jónsson *et al.*, 2013) to check whether this pattern was present in the analysed samples.

### Phylogenetic analyses

To reconstruct the phylogenetic position of the analysed cat remains we used large dataset of sequences of contemporary wildcats and domestic cats published by Driscoll *et al.* (2007). The final dataset consisted of 160 distinct haplotypes encompassing the 2 604 bp long fragment between positions 12 642 and 15 245 of cat’s mtDNA. Phylogenies were reconstructed with Bayesian and Maximum Likelihood methods using MrBayes 3.2.6 (Ronquist *et al.*, 2012) and PhyML 3.1 (Guindon *et al.*, 2010). Best partitioning scheme and substitution model for Bayesian analysis was found with PartitionFinder 2.1.1 (Lanfear *et al.*, 2016) (Supplementary Table 2 & 3). The analysis consisted of two independent runs with four chains each, and was run for 10 000 000 generations with parameters sampled every 1 000 generation. Stationarity and convergence were assessed in Tracer v. 1.6 (ESS>200) (Rambaut and Drummond, 2007). We also confirmed the average standard deviation of split frequencies to be below 0.01. In Maximum Likelihood analysis the HKY + G substitution model was used as indicated by jModeltest 2 (Darriba *et al.*, 2012). The best tree was chosen out of those obtained with NNI and SPR tree rearrangement algorithms, approximate likelihood-ratio test with Shimodaira-Hasegawa ([SH]-aLRT) procedure was applied to assess branch support.

### Radiocarbon dating

Radiocarbon dating of selected samples was performed in Poznan Radiocarbon Laboratory using accelerator mass spectrometry method. Obtained ^14^C dates were calibrated in OxCal v. 4.2.4 (Bronk Ramsey, 2009) using IntCal13 calibration curve (Reimer *et al.*, 2013).

## Results

Out of 20 samples used in hybridization capture experiment 18 produced targeted mtDNA fragment with minimum of 70% sites covered at least two times and those samples were used in phylogenetic reconstruction (Supplementary Table 4). In case of sample Lo1 the recovered fragment of mtDNA was too short to be used in phylogenetic reconstruction but confirmed subspecies assignation. DNA molecules of majority of the samples exhibit damage pattern typical for ancient DNA, only in case of two youngest samples Bis and Ap1 the pattern was questionable (Supplementary Fig. 1). This is however expected, as the amount of damage is the function of time after deposition. Careful examination of bam alignments revealed no signs of contamination. There was also no reads mapping to cat’s mtDNA genome in extraction negative controls.

Reconstructed phylogeny correspond to this obtained earlier by (Driscoll *et al.*, 2007) with clearly separated lineages of European wildcats and Near Eastern wildcats/domestic cats with five sublinages (A – E) distinguished within the latter (Fig. 1). Within sublineage A a branch recently marked A1 by (Ottoni *et al.*, 2017) was observed with moderate support values. Phylogenetic analyses confirmed the initial subspecies assignation and 11 samples were classified as *F. s. lybica/catus* and seven as *F. s. silvestris* (Fig. 1). Out of 11 specimens with *F. s. lybica/catus* mtDNA haplotypes, two specimens yielded modern, two Late Medieval, one Early Medieval and one Roman ages according to radiocarbon dating, while the five other yielded surprisingly early ages of Middle to Late Neolithic, ranging between 5 300 and 4 200 years cal BP (Fig. 2; Supplementary Table 5). The reliability of dating was confirmed by measurements of the C/N ratio in collagen, which was in accepted range (2.9 – 3.6) (DeNiro, 1985). Only in case of one Neolithic sample the collagen yield was too low to confirm quality of the dated material (Supplementary Table 5). Those five samples come from three paleontological sites, Shelter in Krucza Skala (Ks1), Perspektywiczna Cave (Pe1, Pe4, Pe5) and Shelter in Smoleń III (Sh4) and were not associated with cultural remains. Sample Ks1 belonged to sublineage A, Sh4 to A1 while samples Pe1, Pe4 and Pe5 to sublineage B. Samples Pe1, Pe4 and Pe5 yielded similar radiocarbon dates and mtDNA haplotypes, although bones comes from different, non-contiguous layers and distant parts of the site, we cannot exclude possibility that they belong to the single individual. *F. s. lybica/catus* specimens dated to the Roman period until modern times comes both from anthropogenic (Ka1, Sl1, Bis, Bo2) and paleontological (Ap1, Pe8) contexts (Supplementary Table 1). Most of them belonged to mtDNA sublineage C (Ka1, Sl1, Bis, Ap1), while Bo2 belonged to sublineage D and Pe8 to A.

**Figure 1:**
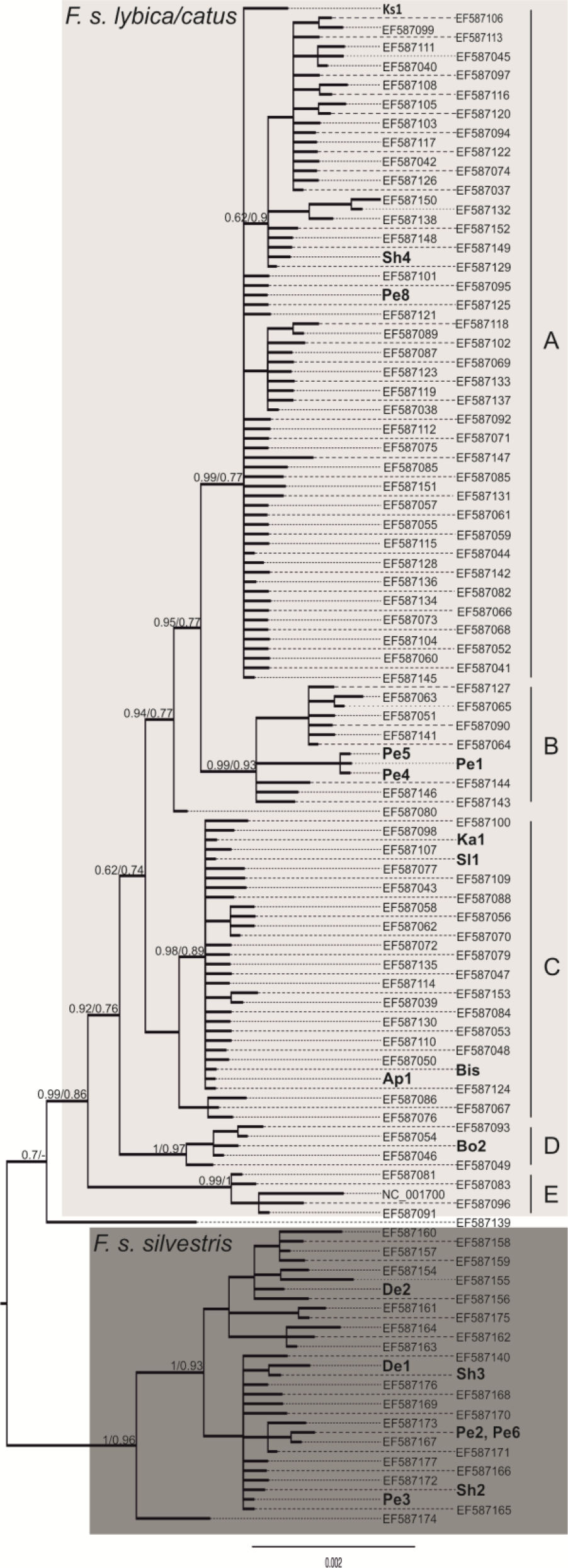
Phylogeny of Holocene and contemporary cats.x. Bayesian phylogeny based on 160 mtDNA haplotypes of Holocene and contemporary cats. Haplotypes of studied samples are bolded. Numbers at nodes indicate posterior probability and SH support values obtained with Bayesian and Maximum Likelihood approaches, respectively. The tree was rooted with sequence of *Felis margarita*(not shown).

**Figure 2.**
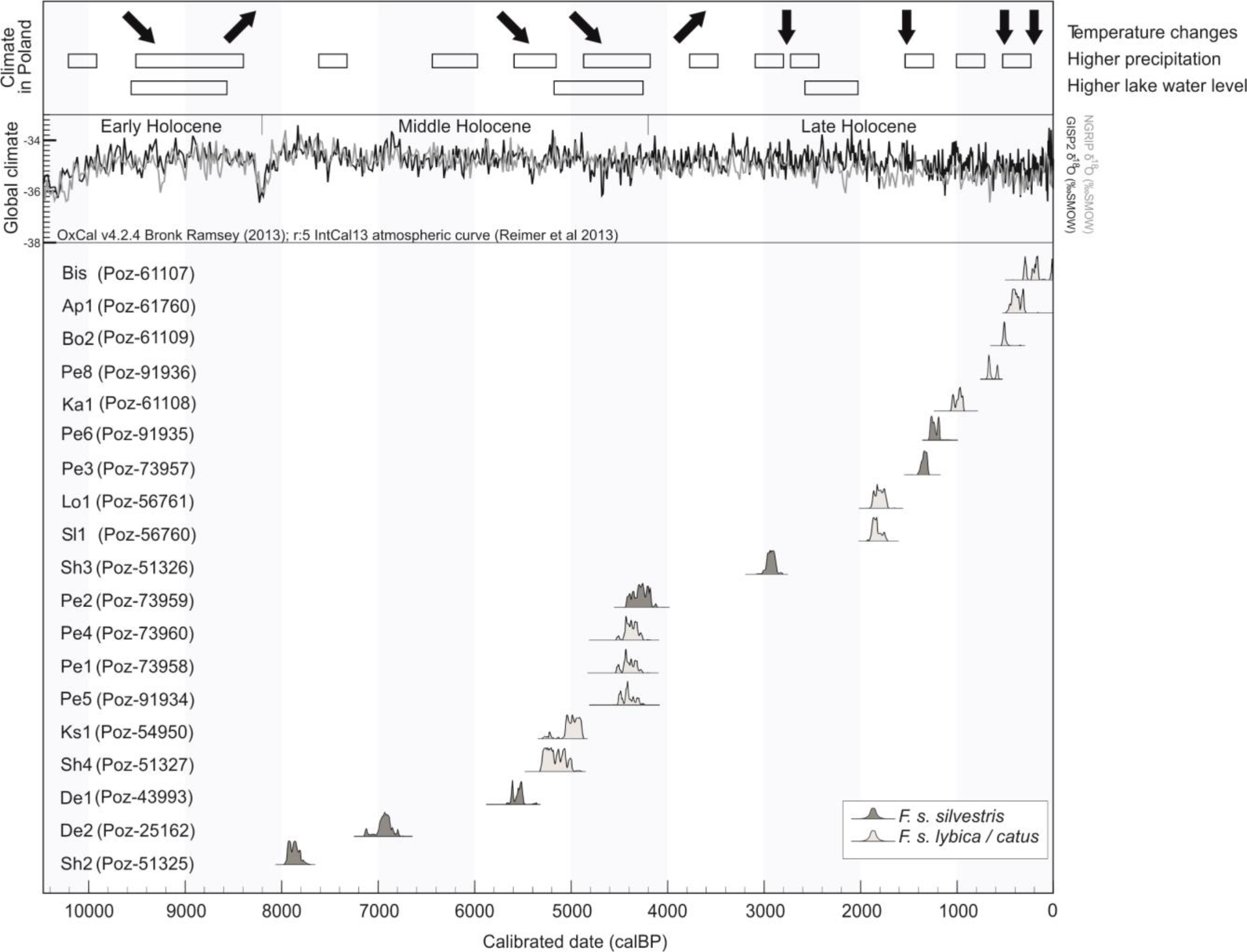
Calibrated radiocarbon ages of the Holocene cat remains. Calibration and δ^18^O curves are given according to (Bronk Ramsey, 2009). The climatic proxies, set in the same scale, are given after (Starkel *et al.*, 2006, 2013).

## Discussion

Available archaeological data suggest that domestic cats made their way to Greece and Rome with Phoenician traders not earlier than 3 400 and 2 500 years BP, respectively (Faure and Kitchener, 2009). Their subsequent spread throughout Europe was mediated by growing Roman Empire and took place around 2 000 years BP, thus the appearance of *F. s. lybica/catus* haplotypes in Poland already in Neolithic period was highly unexpected. Due to the lack of anthropogenic context, the studied *Felis* remains dated to Neolithic period, cannot be easily associated with humans, and other scenarios that led to the ancient occurrence of those haplotypes in Central Europe need to be considered as well. Firstly, their presence could have resulted from natural expansion of *F. s. lybica* from the Near East during the period of favourable climatic conditions. Secondly, it could have resulted from ancient hybridization between European and the Near Eastern wildcats, and subsequent spread of the introgressed individuals into Central Europe. Lastly, the Near Eastern wildcat specimens might have followed humans as synanthropic commensals during the expansion of Neolithic cultures.

The first scenario seems the least likely one. Near Eastern wildcats inhabit mostly hot and dry climatic zones of Northern Africa and Arabian Peninsula with steppe environments including savannas and shrub grasslands (Yamaguchi *et al.*, 2015). Paleoclimate data suggest that the period when the Near Eastern wildcat’s haplotype appeared in Poland was characterized by relatively cool and moist climate with high rate of precipitation and elevated water level (Starkel *et al.*, 2006, 2013) (Fig. 2). This, together with co-occurrence of native European wildcat that mostly inhabits forests, makes the natural expansion of *F. s. lybica* into territory of modern Poland implausible. The second scenario that assumes ancient hybridization of European and the Near Eastern wildcats is more credible. Nowadays hybridization between European wildcats and feral domestic cats is common (Randi *et al.*, 2001; Oliveira *et al.*, 2008; Hertwig *et al.*, 2009; Mattucci *et al.*, 2013). Moreover, Driscoll *et al.*, (2007) found 28 individuals with domestic cat mtDNA among 108 individuals with purely European wildcat nuclear DNA. Such mito-nuclear discordance was interpreted as a result of hybridization event between wildcat subspecies that might have taken place shortly after the domestic cats were brought into the range of European wildcats. Recently, *F. s. lybica* haplotypes were found also in pre-Neolithic Romania those individuals belonged exclusively to mitochondrial lineage A1 (Ottoni *et al.*, 2017). This finding led authors to conclusion that since the beginning of Holocene the natural range of Near Eastern wildcats was wider and included also Southeastern Europe, which became also a historical hybrid zone for European and Near Eastern wildcats. In consequence, the mitochondrial haplotypes of the *F. s. lybica/catus* might have spread in the *F. s. silvestris* populations in Europe. Similar mito-nuclear discordances were observed in other mammalian taxa and interpreted as a result of hybridization after temporary contact between their populations in the past (Alves *et al.*, 2008; Toews and Brelsford, 2012).

Interestingly, the dating of the oldest remains with *F. s. lybica/catus* haplotypes coincides with the appearance of the early farmers in Poland. The earliest Neolithic settlements of Linear Band Pottery culture in Poland appeared around 7 500 years cal BP (Czekaj-Zastawny, 2017). The peak of Neolithic settlement density falls between 5 500 and 4 500 years cal BP in Kuyavia and between 5 000 and 4 000 in Lesser Poland (Timpson *et al.*, 2014). This leads to the third scenario that hypothesizes a spread of the Near Eastern wildcat throughout Europe as a commensal form that followed human groups during the dispersal of Neolithic cultures. The similar way of spread alongside early farmers was recently well documented for early-domesticated pigs (Larson *et al.*, 2007; Ottoni *et al.*, 2013). Processes of wildcat and wild boar domestication have followed the similar, i.e. commensal, pathway (Larson and Fuller, 2014). In its early stages, during Early Holocene, cats and boars had been attracted to human settlements by food wastes and pests and without any deliberate humans activities (Driscoll *et al.*, 2009). Pigs were, however, recognized as a valuable resource and domesticated much earlier than cats, which remained mostly commensal species for next several thousands of years (Larson and Fuller, 2014). The expansion of wildcats to Europe as commensal animals together with early Neolithic groups might have resulted in the observed pattern with Near Eastern wildcat remains found in paleontological contexts not related with humans. Phylogenetic position of *F. s. lybica/catus* individuals from Neolithic Poland strongly supports this scenario, although the presence of lineage A1 may have resulted from introgression of European wildcats with natural population of Near Eastern wildcats in Southeast Europe. However, the presence of lineages A and B cannot be easily explained this way. Lineage A was the main lineage which was domesticated in the Near East and which is the most frequent lineage in recent domestic cats. Individual belonging to this lineage was reported in Early Neolithic Bulgaria around 6 400 years BP, what was also interpreted as a result of human mediated dispersal (Ottoni *et al.*, 2017). Lineage B, the second domesticated lineage was found so far only in Southeast Anatolia, Jordan and Iran. Given that in the dataset by Ottoni *et al.* (2017) there is no a single instance of European wildcat in Anatolia, it’s unlikely that presence of those lineages in Central Europe may have resulted from introgression between *Felis* subspecies. This suggests rather a scenario where the Near Eastern wildcats spread together with early farmers from Anatolia first to Southeast Europe where they crossbreed with local population and acquired lineage A1 and then further northwest to Central Europe. There is also, however, a range of possible intermediate scenarios that cannot be ruled out, such as hybridization between European and Near Eastern wildcats after arrival of early farmers (i.e. haplogroups A and possibly B) to Southeast Europe.

Interesting is the apparent discontinuity between Neolithic and younger samples. Although based on a limited sample size, it suggests that the cats from Neolithic period steam from different source population than domestic cats brought to Central Europe by Romans and that the gene pool of contemporary European domestic cats might have been established from the two different source populations that contributed in the two different periods. This is in line with the findings by Ottoni *et al.* (2017) who showed that cats introduced to Europe during Classical times belonged mostly to lineage C domesticated in Egypt.

Investigation of mtDNA from Holocene *Felis* remains revealed *F. s. lybica/catus* haplotypes present in Central Europe already in Neolithic period. The available data does not allow for certain discrimination between alternatives explaining their presence, however strongly supports dispersal mediated by humans. This transforms current knowledge and poses new questions about the history of domestic cats in Europe. As there is no evidence for domestic cats in archaeological record prior to Roman Period, how and to what extent cats that spread in Europe during Neolithic participated in the genepool of contemporary cats? Further investigation of Holocene and recent cats with a panel of nuclear markers would enable tracing the ancestry of contemporary domestic cats.

## Acknowledgements

We acknowledge Anna Baca for her help in editing and proofreading of the manuscript. Cat remains from Perspektywiczna Cave and Shelter in Smoleń III were collected during field works supported by the National Science Centre, Poland, grant numbers 2011/01/N/HS3/01299 and 2014/15/D/HS3/01302, and radiocarbon dating was supported by the National Science Centre, Poland, grant number 2014/13/D/HS3/03842. MB was supported by the National Science Centre, Poland, grant number 2015/19/D/NZ8/03878.

## Data archiving

Nucleotide sequences reported in this study were deposited in GenBank under accession no. MG813950 - MG813967.

## Author contributions

MB and MK conceived and coordinated the study; MK, MTK, AM, AN, DM provided samples and radiocarbon dating; DP, HP, and MB participated in laboratory work; MB and DP carried out the phylogenetic analyses; MB, MK and MTK wrote the manuscript with significant input from all the authors. All authors gave final approval for publication.

## Supporting Information

**Supplementary Figure 1** Damage patterns and reads length distributions of all *Felis* samples as generated by mapDamage 2 software.

**Supplementary Table S1** Samples analysed in the study.

**Supplementary Table S2** Data blocks used for PartitionFinder analysis.

**Supplementary Table S3** Partitioning scheme and substitution models used in Bayesian analysis.

**Supplementary Table S4** Details of sequencing and consensus calling results.

**Supplementary Table S5** Details of radiocarbon dates used in this study.

## References

Alves PC, Melo-Ferreira J, Freitas H, Boursot P (2008). The ubiquitous mountain hare mitochondria: multiple introgressive hybridization in hares, genus Lepus. Philos Trans R Soc Lond B Biol Sci 363: 2831–2839.

Benecke N (1994). Der Mensch und seine Haustiere. Die Geschichte einer jahrtausendealten Beziehung. Konrad Theiss-Verlag: Stuttgart.

Bronk Ramsey C (2009). Bayesian Analysis of Radiocarbon Dates. Radiocarbon 51: 337–360.

Clutton-Brock J (1999). A Natural History of Domesticated Mammal. Natural History Museum: London.

Czekaj-Zastawny A (2017). The first farmers from the south - Linear Pottery culture. In: Urbanczyk P (ed) The Past Societies. Polish lands from the first evidence of human presence to the Early Middle Ages 2: 5500-2000 BC, Institute of Archaeology and Ehtnology. Polish Academy of Science: Warsaw, p.

Dabney J, Knapp M, Glocke I, Gansauge M-T, Weihmann A, Nickel B, et al. (2013). Complete mitochondrial genome sequence of a Middle Pleistocene cave bear reconstructed from ultrashort DNA fragments. Proc Natl Acad Sci U S A 110: 15758–63.

Darriba D, Taboada GL, Doallo R, Posada D (2012). jModelTest 2: more models, new heuristics and parallel computing. Nat Methods 9: 772–772.

DeNiro M (1985). Postmortem preservation and alteration of in vivo bone collagen isotope ratios in relation to palaeodietary reconstruction. Nature 317: 806–809.

Driscoll CA, Macdonald DW, O’Brien SJ (2009). From wild animals to domestic pets, an evolutionary view of domestication. Proc Natl Acad Sci U S A 106 Suppl: 9971–8.

Driscoll CA, Menotti-Raymond M, Roca AL, Hupe K, Johnson WE, Geffen E, et al. (2007). The Near Eastern origin of cat domestication. Science 317: 519–23.

Faure E, Kitchener AC (2009). An archaeological and historical review of the relationships between felids and people. Anthrozoos 22: 221–238.

Guindon S, Dufayard J-F, Lefort V, Anisimova M, Hordijk W, Gascuel O (2010). New algorithms and methods to estimate maximum-likelihood phylogenies: assessing the performance of PhyML 3.0. Syst Biol 59: 307–21.

Hertwig ST, Schweizer M, Stepanow S, Jungnickel A, Bohle UR, Fischer MS (2009). Regionally high rates of hybridization and introgression in German wildcat populations (Felis silvestris, Carnivora, Felidae). J Zool Syst Evol Res 47: 283–297.

Horn S (2012). Target Enrichment via DNA Hybridization Capture. In: Shapiro B, Hofreiter M (eds) Ancient DNA. Methods and Protocols., Methods in Molecular Biology (Clifton, N.J.). Humana Press: Totowa, NJ Vol 840, pp 189–95.

Jónsson H, Ginolhac A, Schubert M, Johnson PLF, Orlando L (2013). MapDamage2.0: Fast approximate Bayesian estimates of ancient DNA damage parameters. Bioinformatics 29: 1682–1684.

Krajcarz M, Makowiecki D, Krajcarz MT, Masłowska A, Baca M, Panagiotopoulou H, et al. (2016). On the Trail of the Oldest Domestic Cat in Poland. An Insight from Morphometry, Ancient DNA and Radiocarbon Dating. Int J Osteoarchaeol 26: 912–919.

Lanfear R, Frandsen PB, Wright AM, Senfeld T, Calcott B (2016). PartitionFinder 2: New Methods for Selecting Partitioned Models of Evolution for Molecular and Morphological Phylogenetic Analyses. Mol Biol Evol 34: msw260.

Larson G, Cucchi T, Fujita M, Matisoo-smith E, Robins J, Anderson A, et al. (2007). Phylogeny and ancient DNA of Sus provides insights into neolithic expansion in Island Southeast Asia and Oceania. Proc Natl Acad Sci U S A 104: 4834–4839.

Larson G, Fuller DQ (2014). The evolution of animal domestication. Annu Rev Ecol Evol Syst 45: 115–136.

Li H, Durbin R (2010). Fast and accurate long-read alignment with Burrows-Wheeler transform. Bioinformatics 26: 589–95.

Li H, Handsaker B, Wysoker A, Fennell T, Ruan J, Homer N, et al. (2009). The Sequence Alignment/Map format and SAMtools. Bioinformatics 25: 2078–2079.

Lindgreen S (2012). AdapterRemoval: easy cleaning of next-generation sequencing reads. BMC Res Notes 5: 337.

Maricic T, Whitten M, Pääbo S (2010). Multiplexed DNA sequence capture of mitochondrial genomes using PCR products. PLoS One 5: 9–13.

Mattucci F, Oliveira R, Bizzarri L, Vercillo F, Anile S, Ragni B, et al. (2013). Genetic structure of wildcat (Felis silvestris) populations in Italy. Ecol Evol 3: 2443–2458.

Meyer M, Kircher M (2010). Illumina sequencing library preparation for highly multiplexed target capture and sequencing. Cold Spring Harb Protoc 5: t5448.

Milne I, Stephen G, Bayer M, Cock PJA, Pritchard L, Cardle L, et al. (2013). Using Tablet for visual exploration of second-generation sequencing data. Brief Bioinform 14: 193–202.

Oliveira R, Godinho R, Randi E, Ferrand N, Alves PC (2008). Molecular analysis of hybridisation between wild and domestic cats (Felis silvestris) in Portugal: Implications for conservation. Conserv Genet 9: 1–11.

Ottoni C, Girdland Flink L, Evin A, Georg C, De Cupere B, Van Neer W, et al. (2013). Pig domestication and human-mediated dispersal in western eurasia revealed through ancient DNA and geometric morphometrics. Mol Biol Evol 30: 824–832.

Ottoni C, Van Neer W, De Cupere B, Daligault J, Guimaraes S, Peters J, et al. (2017). The palaeogenetics of cat dispersal in the ancient world. Nat Ecol Evol 1: 139.

Rambaut A, Drummond AJ (2007). Tracer v. 1.5.

Randi E, Pierpaoli M, Beaumont M, Ragni B, Sforzi A (2001). Genetic identification of wild and domestic cats (Felis silvestris) and their hybrids using Bayesian clustering methods. Mol Biol Evol18: 1679–1693.

Reimer PJ, Bard E, Bayliss A, Beck JW, Blackwell PG, Bronk C, et al. (2013). Intcal13 and marine13 radiocarbon age calibration curves 0 – 50,000 years cal bp. Radiocarbon 55: 1869–1887.

Rohland N, Harney E, Mallick S, Nordenfelt S, Reich D (2014). Partial uracil-DNA-glycosylase treatment for screening of ancient DNA. Philos Trans R Soc B Biol Sci 370: 20130624–20130624.

Ronquist F, Teslenko M, van der Mark P, Ayres DL, Darling A, Höhna S, et al. (2012). MrBayes 3.2: efficient Bayesian phylogenetic inference and model choice across a large model space. Syst Biol 61: 539–42.

Starkel L, Michczynska DJ, Krąpiec M, Margielewski W, Nalepka D, Pazdur A (2013). Progress in the Holocene chrono-climatostratigraphy of Polish territory. Geochronometria 40: 1–21.

Starkel L, Soja R, Michczyńska DJ (2006). Past hydrological events reflected in Holocene history of Polish rivers. Catena 66: 24–33.

Timpson A, Colledge S, Crema E, Edinborough K, Kerig T, Manning K, et al. (2014). Reconstructing regional population fluctuations in the European Neolithic using radiocarbon dates: A new case-study using an improved method. J Archaeol Sci 52: 549–557.

Toews DPL, Brelsford A (2012). The biogeography of mitochondrial and nuclear discordance in animals. Mol Ecol 21: 3907–3930.

Vigne J-D, Guilaine J, Debue K, Haye L, Gerard P (2004). Early taming of the cat in Cyprus. Science (80-) 304: 259.

Yamaguchi N, Kitchener AC, Driscoll CA, Nussberger B (2015). Felis silvestris. The IUCN Red List of Threatened Species. Felis silvestris IUCN Red List Threat Species.

